# How sparse can a fly-brain model be? Connection strength, wiring placement, network state and task jointly determine function-preserving compression of the *Drosophila* connectome

**DOI:** 10.64898/2026.09.19.752860

**Authors:** Donggyu An

## Abstract

Whole-brain connectomes are increasingly used as wiring diagrams for spiking simulations and as structural priors for artificial networks, usually after an ad hoc synapse-count threshold whose effect on validated function has never been measured. We define *function-preserving sparsification*—removing edges under a fixed budget while preserving validated behaviour and population responses—and study it in the leaky integrate-and-fire whole-brain model of Shiu et al. (2024). Six operators—weak-synapse pruning, uniform and weight-proportional random removal, degree-preserving rewiring, weight shuffling and a task-aware criterion—were compared at identical edge budgets across three validated circuits (sugar-evoked proboscis extension, its bitter suppression, grooming pathway specificity), silent and spontaneously active network states, and multiple seeds. Weak-synapse pruning preserves 79% of feeding output with 6% of edges, with a cliff at k* ≈ 9 synapses, whereas random, degree-preserving and weight-shuffled controls at the same budget abolish output, and weight-proportional random sampling survives only while it retains the strong-edge set: function resides in the conjunction of wiring placement and strength, not in degree statistics or synaptic mass. The cliff is task-dependent—grooming specificity collapses below k = 2 through disinhibition. In the active state failure changes character: output over-excites instead of vanishing, and bitter suppression is lost progressively from k = 5 and reverses at k = 31. A task-aware criterion preserves its training stimulus at 0.4% of edges but loses held-out inhibition first. These results provide a first benchmark for connectome sparsification and show that what must be kept depends on task and network state.

**Author summary:** Neuroscientists now have a map of every neuron and connection in the fruit-fly brain. Computer scientists have begun to use that map directly—as a circuit to simulate, or as a fixed "wiring diagram" inside artificial networks that steer robots or move simulated bodies. But the map is large—15 million connections—and almost every study quietly discards part of it before use, typically by deleting connections with only a few synapses. Nobody had checked what that deletion does to the behaviours the model is supposed to reproduce. We asked how much of the fly connectome can be removed from an experimentally validated brain simulation before it stops working, and what decides which connections matter. The answer was not simply "keep the strong ones": choosing the same number of connections at random, rewiring them while preserving each neuron’s connection count, or keeping their positions but shuffling their strengths all destroy behaviour. Which circuits survive also differed—a grooming circuit that must distinguish two sensory pathways collapsed much earlier than a simple feeding circuit, and when the brain was spontaneously active, inhibition was the first thing to go. The results give principled grounds for the thresholds connectome-based models use and provide a test bed for future compression methods.

## 1. Introduction

### The connectome has become a computational substrate

The synapse-resolution whole-brain connectome of an adult female *Drosophila melanogaster* (FlyWire) comprises 139,255 neurons and about 50 million synapses, which condense into 15.1 million weighted directed connections between neuron pairs (Dorkenwald et al., 2024; Schlegel et al., 2024). It is the third nervous system to be reconstructed at synapse resolution in its entirety, after *Caenorhabditis elegans* (Cook et al., 2019) and the *Drosophila* larva (Winding et al., 2023), and it completes the earlier partial reconstruction of the adult central brain (Scheffer et al., 2020); the programme all of them belong to was set out two decades ago (Sporns et al., 2005). The connectome has moved beyond anatomy into a computational object. Shiu et al. (2024) built a whole-brain leaky integrate-and-fire (LIF) model constrained only by the connectome and predicted neurotransmitter signs, and confirmed 91% of 164 predictions about sensorimotor circuits experimentally. Lappalainen et al. (2024) showed that a connectome-constrained network with only a few hundred free parameters predicts neural responses across the fly visual system. In 2026 the full connectome graph was used as the recurrent controller of a navigating robot (FLYNN; B. Wang & Chen, 2026) and for whole-body locomotion of a biomechanical fly (FlyGM; Jin et al., 2026), and the whole model was mapped onto neuromorphic hardware (F. Wang et al., 2025), on processors whose energy advantage scales directly with how few synapses have to be represented (Davies et al., 2018; Roy et al., 2019).

### An unexamined sparsification step

All of these uses pay a computational cost proportional to the number of edges, and most reduce it with a threshold. FLYNN keeps only connections with at least five synapses (15.1 M → 5.34 M on FlyWire v783), and Pospisil et al. (2024) apply the same threshold before linear analysis (in the v630 graph used here the same criterion keeps 17.8% of edges, 2.61 M; the difference reflects connectome version and neuron inclusion criteria). FlyGM reports per-step memory and time above those of a multilayer perceptron as a limitation. These thresholds are inherited from anatomical practice—connections of 1–2 synapses are reproduced between hemispheres only 42–60% of the time, whereas connections of ≥ 10 synapses are preserved above 90% (Schlegel et al., 2024)—but reproducibility *between* brains is not the same as functional necessity *within* a brain. No study has measured how a model’s validated functions change as a function of sparsification budget, or compared the threshold heuristic with alternatives at the same budget.

### A parallel problem in artificial networks

Removing parameters from a trained artificial network is an old problem: second-derivative saliency (LeCun et al., 1989; Hassibi et al., 1993), magnitude pruning with retraining (Han et al., 2015), and the finding that sparse subnetworks trained from their original initialisation can match the dense network (Frankle & Carbin, 2019). The field has since divided into rules that score connections before training (Lee et al., 2019; C. Wang et al., 2020), rules that maintain sparsity throughout training (Mocanu et al., 2018; Evci et al., 2020), and rules that learn the mask while fine-tuning (Sanh et al., 2020); Hoefler et al. (2021) survey the resulting landscape. What almost all of them share is a recovery step: once edges are removed the surviving weights are retrained or rewound (Renda et al., 2020), and careful evaluations have found that the accuracy reached afterwards depends at least as much on the surviving architecture and the training budget as on the saliency rule that selected it (Gale et al., 2019; Liu et al., 2019). A meta-analysis of 81 papers identified the central weakness of that literature as the absence of standardised benchmarks and controls, which makes competing pruning rules hard to compare (Blalock et al., 2020). The connectome setting differs in two ways that make it informative for both fields. First, the network is not trained: weights are fixed measurements, so a removed edge cannot be compensated by retraining and the contribution of wiring is isolated rather than confounded with learning. Second, the target of preservation is not one accuracy number but a set of experimentally validated input–output behaviours together with the population response that produces them, so a rule can be scored on several axes at once. What follows supplies for a connectome what Blalock et al. asked for in artificial networks: a fixed budget, one teacher, matched null models and several preserved quantities.

### Why the answer is not obvious

Two lines of evidence suggest that the answer depends on what is controlled. Dhiman (2026) showed that the reported advantage of connectome topology in trained networks disappears when initialisation is shared and a degree-preserving null model is used, whereas FlyGM found a 1.6-fold advantage on a harder locomotion task even against the same degree-preserving control (Jin et al., 2026). In an LIF model every neuron is silent at rest, so weak synapses onto silent neurons do nothing; in a spontaneously active network, thousands of weak synapses summing near threshold may matter. Whether the functionally necessary subgraph is a fixed property of the wiring or a function of network state and task is an open question, and it determines which sparsification results transfer between models.

### What a null model has to control

A claim that a measured network differs from chance is only as strong as the ensemble it is compared against, and network neuroscience has converged on a family of randomisations that each destroy one property while preserving others: degree-preserving edge swapping (Maslov & Sneppen, 2002; Rubinov & Sporns, 2010), reshuffling of weights over a fixed topology, and strength-or geometry-matched ensembles (Betzel & Bassett, 2018). Váša and Mišić (2022) review these choices and show how strongly a conclusion can turn on which one is made; the same concern has now reached connectome-constrained modelling (Dhiman, 2026). Because the quantity to be preserved here is not a graph statistic but a simulated behaviour, we adopt that logic directly: every operator is applied at an identical edge budget, and the nulls are chosen so that degree sequence, placement and strength are each removed on their own. The question they answer is a biological one as well, since nervous systems are themselves thought to trade wiring cost against function (Bullmore & Sporns, 2012), which makes the functional value of the weakest connections worth measuring rather than assuming.

### This study

We formalise function-preserving sparsification of a connectome as a constrained subgraph-selection problem (Methods) and study it in the LIF model of Shiu et al., the only whole-brain model whose function has been validated experimentally. Six sparsification operators are compared at the same edge budget across three validated circuits—sugar-evoked proboscis extension and its bitter suppression, whose sensory and premotor components have been characterised genetically (Sterne et al., 2021; Shiu et al., 2022), and mechanosensory grooming, whose command pathway and suppression hierarchy are known (Seeds et al., 2014; Hampel et al., 2015)—silent and spontaneously active states, degree-preserving, weight-shuffled and weight-proportional random null models, and multiple seeds. We report (i) budget–function curves and the position of the functional cliff, (ii) which graph properties are required, (iii) whether behavioural and neural-response preservation coincide, and (iv) how all of this changes with state and task. The central finding is that **the limit of function-preserving compression is not a fixed threshold on connection strength but is set jointly by the conjunction of strength and placement, by the inhibition a task requires, and by network state.**

## 2. Results

### 2.1 The sparsification problem and the reference model

The reference graph *G=(V,E,w)* is FlyWire materialisation 630 as used by Shiu et al. (127,400 of the 139,255 FlyWire neurons that Shiu et al. included in the simulation; 14,687,178 edges; *w_ij* = signed synapse count, sign from predicted neurotransmitter). A sparsification operator returns *G’=(V,E’,w|_E’)* with *E’⊂ E*, *|E’|=b*. Function is measured by simulating validated stimulus protocols (Poisson input to defined sensory populations) in *G* and *G’*: (a) *behavioural readout*—the firing rate of the validated output neuron (MN9 for the feeding circuit, aBN1 for grooming) as a ratio to the full model; (b) *neural response*—the Pearson correlation of per-neuron mean rates over the union of neurons active in either model, and the Jaccard overlap of the active sets (stimulated neurons excluded; Methods). All operators are evaluated at the same budget *b_k = |{(i,j): |w_ij|>k}|*, *k∈{1,2,3,5,10,20,31,50}*, so differences between operators reflect *which* edges are kept, not how many. Because the budget *k* means "more than *k* synapses" (i.e. at least *k+1*), the anatomical thresholds "≥ 5" and "≥ 10" correspond to *k = 4* and *k = 9* on our grid (17.8% and 7.0% of edges; *k = 5* and *10* keep 14.1% and 6.1%). *k = 10* and *k = 31* are the grid points closest to the 90% and 99% inter-hemispheric reproducibility thresholds reported by Schlegel et al. (≥ 10 and ≥ 31 synapses).

### 2.2 Weak-synapse pruning preserves behaviour beyond the conventional threshold (≥ 5), up to the inter-hemispheric reproducibility threshold (≥ 10) (H1)

In the silent (σ = 0) sugar circuit, removing every single-synapse edge (50% of edges, 14% of synapse mass) leaves the MN9 response unchanged (ratio 1.04, population correlation 0.997; Figs 1b and 2a). At k = 5 (14% of edges, 57% of mass) the ratio is 0.95 and the correlation 0.95; at k = 10 (6% of edges, 41% of mass) 0.79 and 0.82; the ratio then drops to 0.11 at k = 20, 0.09 at k = 31 and 0 at k = 50. The cliff position, log-interpolated at a ratio of 0.8, is **k* ≈ 9.4** (≈ 10 synapses), close to the 90% inter-hemispheric reproducibility threshold (≥ 10 synapses). The number of active neurons fell faster than the readout (361 → 322 at k = 5 → 269 at k = 10 → 132 at k = 20). The sugar-only condition of the bitter task reproduced the same curve (1.09, 1.00, 0.86, 0.10 at k = 2, 5, 10, 31).

**Figure 1.**
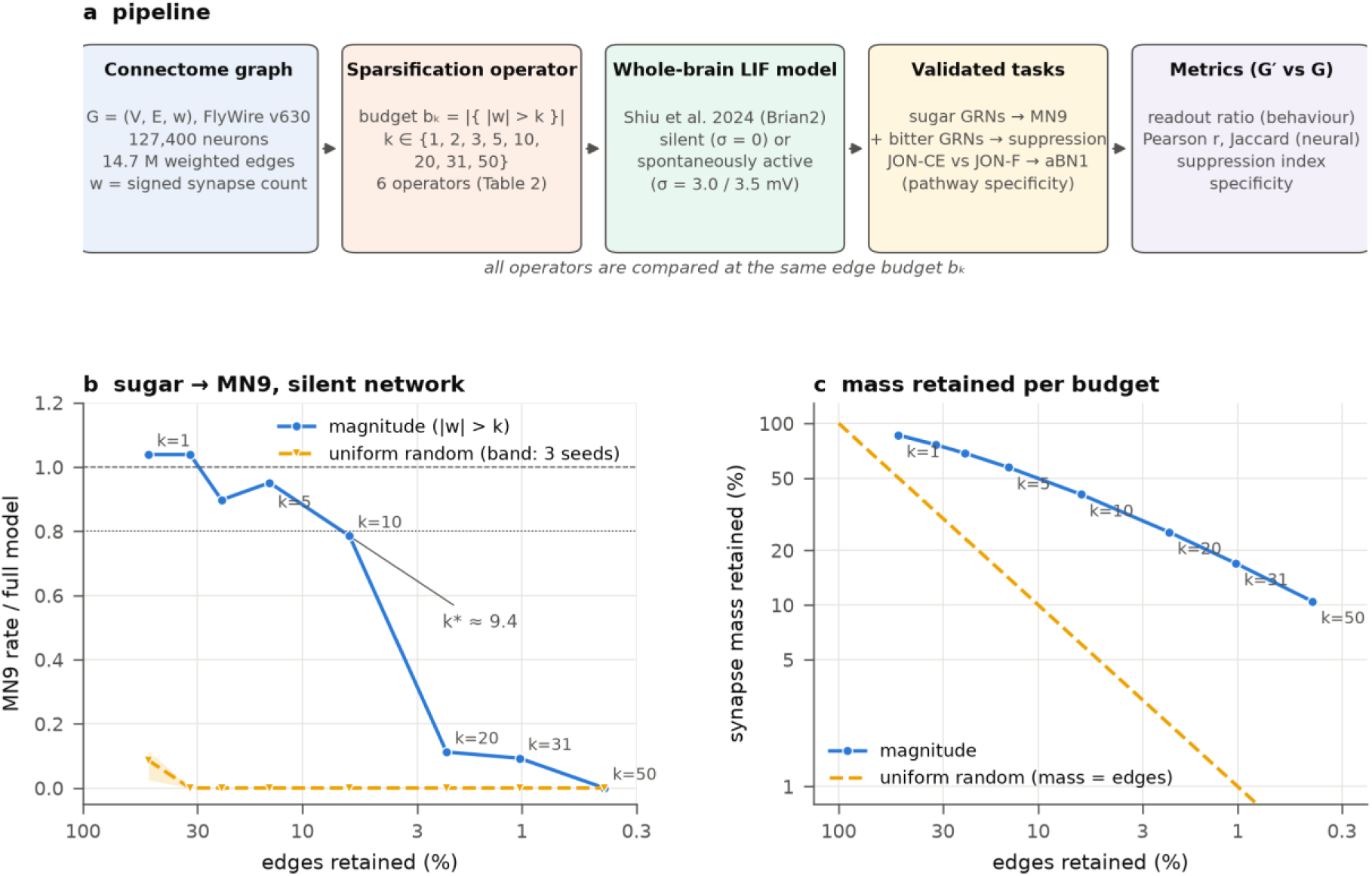
Function-preserving sparsification of a validated whole-brain model. (a) Pipeline. Six sparsification operators (Table 2) are applied to the FlyWire v630 connectome (127,400 neurons, 14.7 M weighted edges) at the same edge budget *b_k = |{|w_ij| > k}|*; three validated tasks are simulated in the LIF whole-brain model of Shiu et al. (2024) in the silent (σ = 0) or spontaneously active (σ = 3.0, 3.5 mV) state and compared with the full model. (b) Sugar → MN9 readout ratio in the silent state. Magnitude pruning (blue) holds 0.79 up to k = 10 (6% of edges) with a cliff at k* ≈ 9.4; a uniform random subset at the same budget (yellow; band = range over 3 seeds) gives 0.09 at k = 1 and 0 thereafter. (c) Synapse mass retained at each budget. Weak-synapse pruning keeps 41% of the mass with 6% of the edges, whereas uniform random keeps mass in proportion to edges (dashed). In all figures the x-axis "edges retained (%)" is logarithmic and sparser to the right.

**Figure 2.**
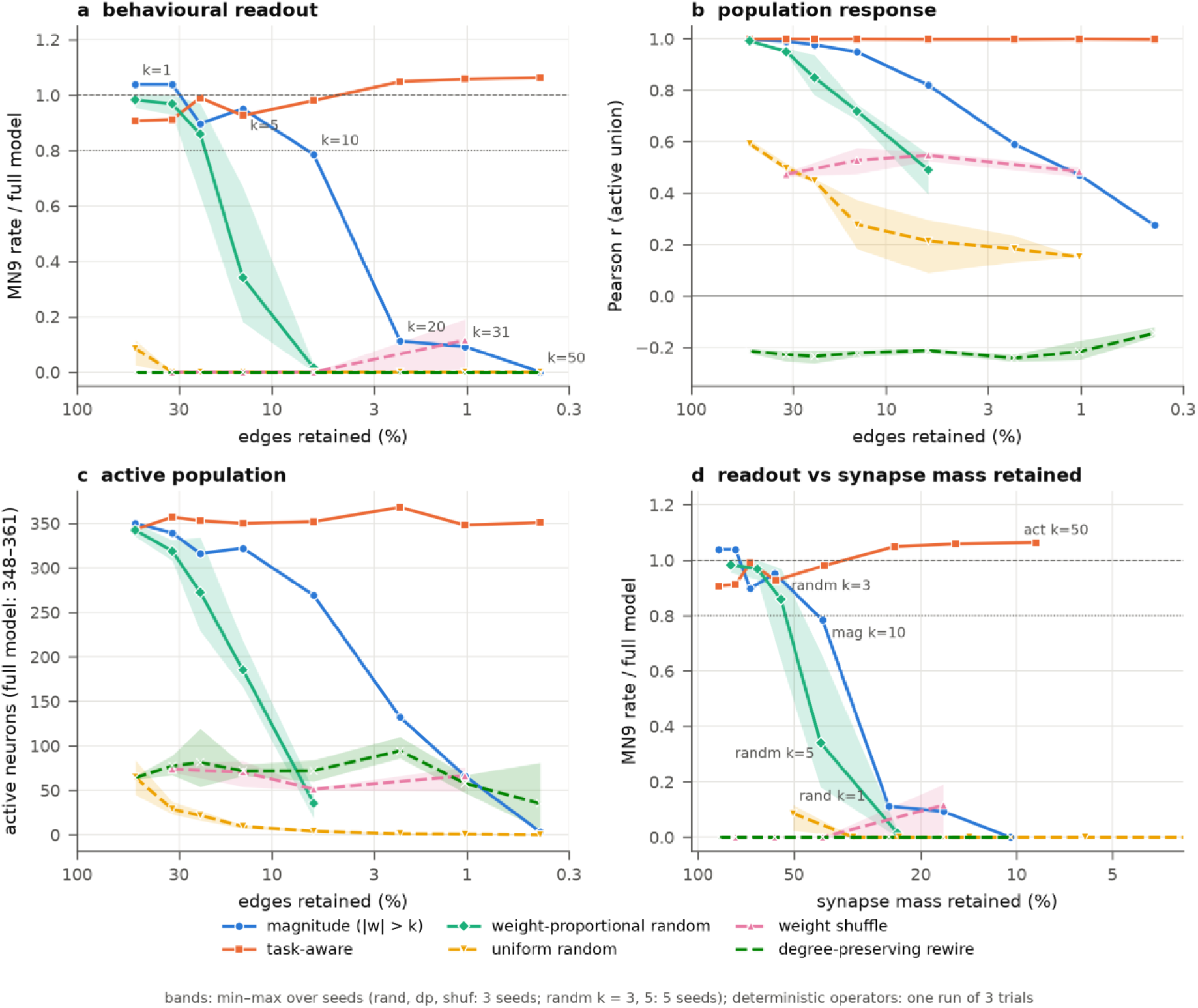
Which edges are kept decides survival (sugar → MN9, σ = 0). (a) Readout ratio, (b) population Pearson correlation over the union of active neurons, and (c) number of active neurons (full model 348– 361) against the fraction of edges retained. Solid lines: magnitude (mag), task-aware (act), weight-proportional random (randm); dashed: uniform random (rand), weight shuffle (shuf), degree-preserving rewiring (dp). Bands are seed ranges of the random operators (rand, dp, shuf: 3 seeds; randm at k = 3, 5: 5 seeds). Numbers give k. (d) The readout ratio replotted against synapse mass retained. randm k = 5 and mag k = 10 (41%), and rand k = 1 and randm k = 3 (≈ 50%), have equal mass but opposite outcomes, so total synapse mass does not explain survival.

**Table 1.** Budget *k* and the retained graph.

| $k$ | edges retained | synapse mass retained | anatomical meaning |
| --- | --- | --- | --- |
| 1 | 50.3% | 86.2% | single-synapse connections removed |
| 2 | 32.5% | 76.3% |  |
| 3 | 23.3% | 68.6% |  |
| 5 | 14.1% | 57.3% | ≈ FLYNN / Pospisil threshold ($\geq 5$ synapses = 17.8%) |
| 10 | 6.1% | 40.7% | $\approx 90\%$ inter-hemispheric reproducibility ($\geq 10$ synapses = 7.0%) |
| 20 | 2.2% | 25.2% |  |
| 31 | 1.0% | 17.0% | $\approx 99\%$ inter-hemispheric reproducibility |
| 50 | 0.4% | 10.5% |  |

**Table 2.**
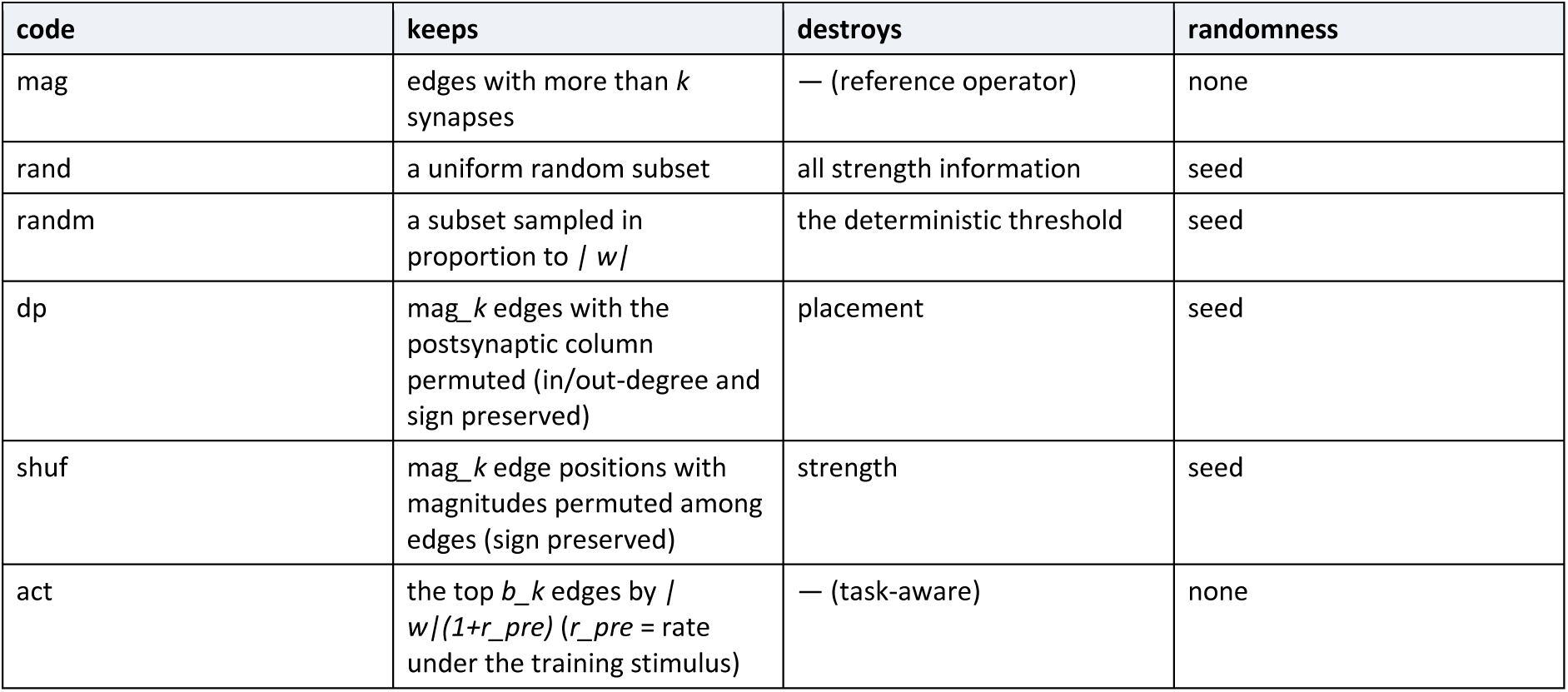
Sparsification operators. (all at the same budget *b_k*; definitions in Methods 4.3)

### 2.3 Neither degree statistics, nor placement alone, nor strength alone, nor synaptic mass alone is sufficient (H3)

Three controls with the same number of edges failed at every budget (Fig. 2). A **uniform random subset** (rand) already reduced the MN9 ratio to 0.02–0.12 at k = 1 (3 seeds) and to 0 for k ≥ 2. **Degree-preserving rewiring** (dp)—permuting the postsynaptic column of the edges retained by weak-synapse pruning, which preserves every neuron’s in-and out-degree and Dale’s sign exactly—gave a ratio of 0.00 at every k and seed, with *negative* population correlations (−0.14 to −0.24): the rewired network activates a different population. dp was 0 without exception in the bitter and grooming tasks as well (Phase B, all 48 conditions). **Weight shuffling** (shuf)—permuting synapse-count magnitudes among the same edge positions, signs preserved—gave a ratio of 0.00 at k = 2, 5 and 10 in the bitter task, reduced the active population from 355 to 51–82 neurons, and gave population correlations of 0.47–0.58; only at k = 31 did it partially recover (0.02–0.17), because at that budget every retained edge has more than 31 synapses and permutation barely changes the magnitude distribution. In the grooming circuit shuf failed differently: the JON-CE response of aBN1 fluctuated between 0.03 and 2.38 across seeds (k = 2–10), the JON-F response—0.33 Hz in the full model—appeared at 0–24 Hz depending on the seed, and pathway specificity became **a random value decided by the seed** (1.7–39.7 Hz at k = 2, −16.3–21.3 Hz at k = 5, −17.0–40.3 Hz at k = 10; full model 24.0), with population correlations of 0.30–0.55. Preserving placement while losing magnitude thus leaves it to the accident of the permutation which pathway drives the readout neuron. shuf behaved the same in the sugar task: at k = 2, 5 and 10 all three seeds gave an MN9 ratio of 0.00, active neurons 348 → 50–83, population correlation 0.46–0.58, and only k = 31 gave 0.01–0.19 (mean 0.12). In the spontaneously active state (σ = 3.5) the sugar response of shuf in the bitter task was 0.01–0.30. The failure of shuf is therefore independent of task and state.

Do random controls die simply because they lose synaptic *mass* (the sum of retained synapse counts)? To test this objection we added a **weight-proportional random** operator (randm) that draws the same number of edges without replacement with probability proportional to *|w_ij|* (Fig. 2d). randm retains far more mass than uniform random: 79, 65, 55, 41 and 24% of the total at k = 1, 2, 3, 5 and 10 (mag: 86, 76, 69, 57, 41%; rand: 50, 32, 23, 14, 6%). It includes 99–100% of edges with more than 31 synapses for k ≤ 3, and 99, 93 and 83% of edges with more than 10 synapses at k = 1, 2 and 3. Accordingly randm was indistinguishable from magnitude pruning at k = 1 and 2 (MN9 ratio 0.96–1.01 and 0.93–1.00 versus 1.04, 1.04), and at k = 3 (5 seeds) its mean of 0.86 matched magnitude pruning (0.90) although seed-to-seed spread began to appear (0.64–0.97). At k = 5 (5 seeds), however, it collapsed to 0.18, 0.20, 0.67, 0.26 and 0.40 (mean 0.34; mag 0.95), and at k = 10 to 0.00–0.03 (mag 0.79)—exactly where randm retains only 63% (k = 5) and 34% (k = 10) of the edges with more than 10 synapses. Population correlation and active-population size departed before behaviour did (at k = 3: correlation 0.78–0.94, 229–334 active neurons; mag 0.98, 316). Comparison at equal mass makes the point: randm k = 5 (0.34) versus mag k = 10 (0.79) at 41% of mass; randm k = 10 (0.02) versus mag k = 20 (0.11) at 24%; and conversely rand k = 1 (50% of mass, ratio 0.09) versus randm k = 3 (55%, 0.90) are opposite outcomes at similar mass. The failure of random controls therefore does not reduce to mass loss. What decides survival is not the total retained synapse count but **how completely a specific set of strong edges is retained**, and randm collapses exactly at the budget where it begins to miss that set stochastically.

Functional information is thus carried by edge placement and edge strength **together**. The degree sequence (the null of Dhiman, 2026), placement alone, strength alone and synaptic mass alone all fail to recover it—the "dp dies + shuf dies → both placement and strength are needed" row of the pre-registered interpretation table (Design §3 H3). That shuf retains a higher population correlation than dp (0.5 versus −0.2) shows that placement carries part of *which neurons are active*, but the few strong edges needed to reach the validated output fall below threshold once they are given typical (median 2– 3) magnitudes.

### 2.4 The task-aware criterion wins on its training stimulus but loses inhibition (H6)

Ranking edges by *|w_ij|(1+r_i)*—where *r_i* is the presynaptic firing rate in the full model under the training stimulus (sugar100)—(act) preserved the MN9 ratio at 0.91–1.06 for every k at σ = 0 (Fig. 2), with population correlation ≥ 0.997 and 343–368 active neurons (full model 361): **both behaviour and neural response to the training stimulus survive at 0.4% of edges (k = 50)**, in contrast to magnitude pruning, which collapses to 0.11 at k = 20.

Functions not used in training behaved differently (Fig. 5c, d). In the bitter task act kept the sugar-only response (1.09, 0.95, 1.00, 0.96 at k = 2, 5, 10, 31) while raising MN9 under sugar+bitter from 2.0 Hz in the full model to 4.0, 8.7, 16.0 and **68.7 Hz**. The suppression index fell from 0.97 (full) to 0.95, 0.86, 0.76 and **−0.06**, below the magnitude criterion at the same budget (0.96, 0.87, 0.93, 0.32) from k = 10 onwards. The activity-weighted score assigns low scores to the inhibitory pathways that are silent under the training stimulus and removes them first, yielding a graph that is perfect on the training stimulus but "disinhibited" and unresponsive to bitter. The same held in the active state (σ = 3.5): act’s sugar response stayed at 1.0–1.46, but its suppression index at k = 2, 5, 10, 31 was 0.91, 0.80, −0.29, 0.13 (3 trials), collapsing at k = 5–10 together with the magnitude criterion (10 trials: 0.91, 0.72, 0.40, −0.61).

On the held-out grooming stimulus (JON-F) transfer succeeded. act trained on jonCE100 kept the JON-CE response at 1.00–1.14 and population correlation ≥ 0.99 in the silent state while holding the JON-F response at 0–1.7 Hz (10.3 Hz only at k = 5), preserving pathway specificity of 24.0 Hz at 26.7 Hz even at k = 31; at σ = 3.5 it kept the JON-CE response at 0.95–0.98 and specificity at 21.3 → 22.7–35.7 Hz for k = 2, 5, 10 (although at k = 31 the training-stimulus response itself fell to 0.42, specificity 15.3 Hz—in the active state, act’s preservation of its own training stimulus also reaches its limit at 1% of edges). The difference between the two tasks lies in what the held-out function requires. Bitter suppression needs an *additional* pathway that is silent under the training stimulus (bitter GRNs → inhibitory neurons) to remain in the graph, whereas grooming specificity only requires that the JON-F pathway *not* drive aBN1, so act’s bias against edges silent under training happens to coincide with the correct answer. Task-specific compression thus preserves "the absence of other functions" for free but loses "the presence of other functions". This is both a methodological warning against reusing a compressed graph built from one stimulus for other functions, and a criterion predicting where transfer will fail.

### 2.5 The cliff position depends on the task: pathway specificity collapses first (H4)

The grooming circuit consists of two sensory pathways with similar synapse counts but different functions (JON-CE → aBN1 evokes grooming; JON-F → aBN1 does not), and so tests "placement rather than strength" most directly (Fig. 5a, b). In the silent state magnitude pruning reduced the JON-CE response to 0.73 already at k = 2, 0.23 at k = 5 and 0.03 at k = 10—far earlier than the sugar circuit (1.04, 0.95, 0.79 at the same k). At the same time the JON-F response, 0.33 Hz in the full model, **increased** to 5.0 Hz at k = 2 and 9.0 Hz at k = 5, so that pathway specificity (aBN1_CE − aBN1_F) collapsed from 26.7 Hz to 14.7 (k = 2), −2.7 (k = 5) and 0.3 (k = 10). The grooming cliff is therefore at **k* < 2**. The anatomical analysis (§2.8) shows why: the input that the intermediates of the JON-F → aBN1 pathway give to aBN1 is inhibition-dominated (222 excitatory versus 671 inhibitory synapses), so when weak inhibitory edges go first, the normally suppressed F pathway is disinhibited.

Bitter suppression was more robust. In the silent state the suppression index SI = 1 − MN9(sugar+bitter)/MN9(sugar) stayed at 0.96, 0.87 and 0.93 for k = 2, 5, 10 (full model 0.96) before dropping to 0.32 at k = 31 (where the sugar response itself has collapsed to 0.10, making SI unstable). The rise of sugar+bitter MN9 from 2.3 to 8.7 Hz at k = 5 reproduces the partial disinhibition seen in the pilot. In the silent state the three cliffs are therefore ordered **grooming specificity (k *< 2) ≪ sugar output (k* ≈ 9) ≲ bitter suppression (10 < k* < 31)**, and evaluating compression by a single output misses the loss of circuit function.

### 2.6 In the spontaneously active state the mode of failure changes: over-excitation and loss of inhibition rather than silence (H2)

With membrane noise σ = 3.5 mV the full model is in a state where 78–93% of neurons are active and the mean spontaneous rate is 4.3 Hz (Methods; the fraction depends on stimulation; Figs 3 and 4). In this state the sugar-evoked MN9 rate itself is highly variable across trials (3-trial mean 49.3 ± 36.1 Hz, 10-trial mean 37.0 ± 25.5 Hz; silent state 68.3 ± 3.6), so a cliff is hard to define from the readout ratio alone. With that caveat, pre-registered prediction A (an earlier cliff) was not supported. In the 10-trial run the magnitude-pruning readout ratio was **≥ 1.03** at k = 1, 3, 5, 10, 20 and 31 (1.03, 1.03, 1.59, 1.23, 1.19, 1.45; absolute rates 38–59 Hz against a full-model 37.0 ± 8.1 Hz, mean ± SE), and at σ = 3.0 it was 1.22–1.64 for k = 1–20. The 3-trial run estimated a higher reference (49.3 Hz) and hence ratios of 0.59– 1.24, but the absolute rates (29–61 Hz) were in the same range. The output silencing that appears at k = 20 in the silent state therefore does not appear up to k = 31 in the active state; output is at or above the full-model level. Population correlation declined gradually (σ = 3.5: 0.99, 0.94, 0.88, 0.79, 0.70, 0.60 at k = 1, 5, 10, 20, 31, 50; k = 50 from the 3-trial run) and the σ = 3.0 curve (0.99, —, 0.88, 0.79, 0.71) was nearly identical, so the result is robust to noise amplitude. Compared with the silent-state correlation curve (0.997, 0.95, 0.82, 0.59, 0.47, 0.27) it is in fact higher for k ≥ 20, because the fan-in structure of the 100,000 noise-activated neurons sustains the correlation.

**Figure 3.**
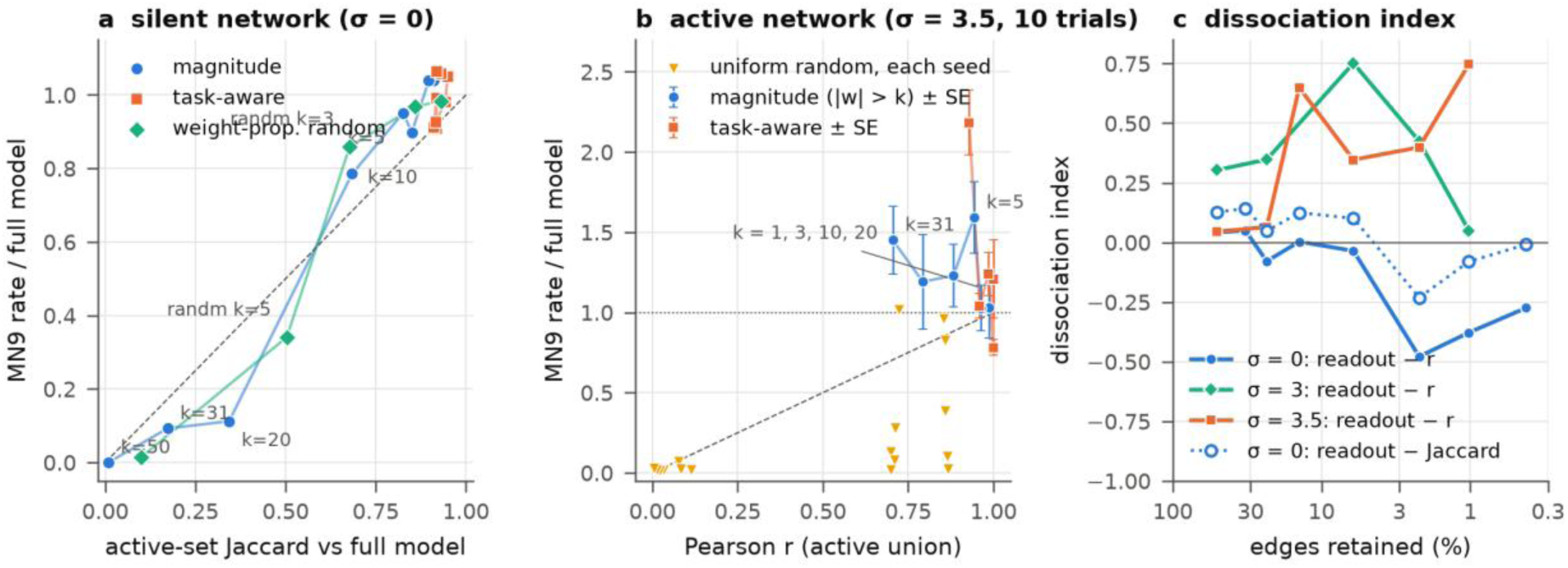
Behavioural output and neural response dissociate. (a) Silent state: readout ratio (y) against active-set Jaccard (x) for magnitude, task-aware and weight-proportional random (points = seed means per k). Above the diagonal, behaviour holds while the active set changes; magnitude pruning is in this region at k = 5–10. (b) Active state (σ = 3.5, 10 trials): readout ratio (± trial SE) against population Pearson correlation. Magnitude pruning exceeds the full model while the correlation falls to 0.7; uniform random (yellow, per seed) splits between 0.02 and 0.96 at a correlation of 0.85. (c) Dissociation index (readout ratio − population correlation) against budget by state; dotted line, Jaccard-based index at σ = 0. Negative for k ≥ 20 in the silent state (behaviour goes first), positive at every k in the active state (behaviour outlives the neural response).

**Figure 4.**
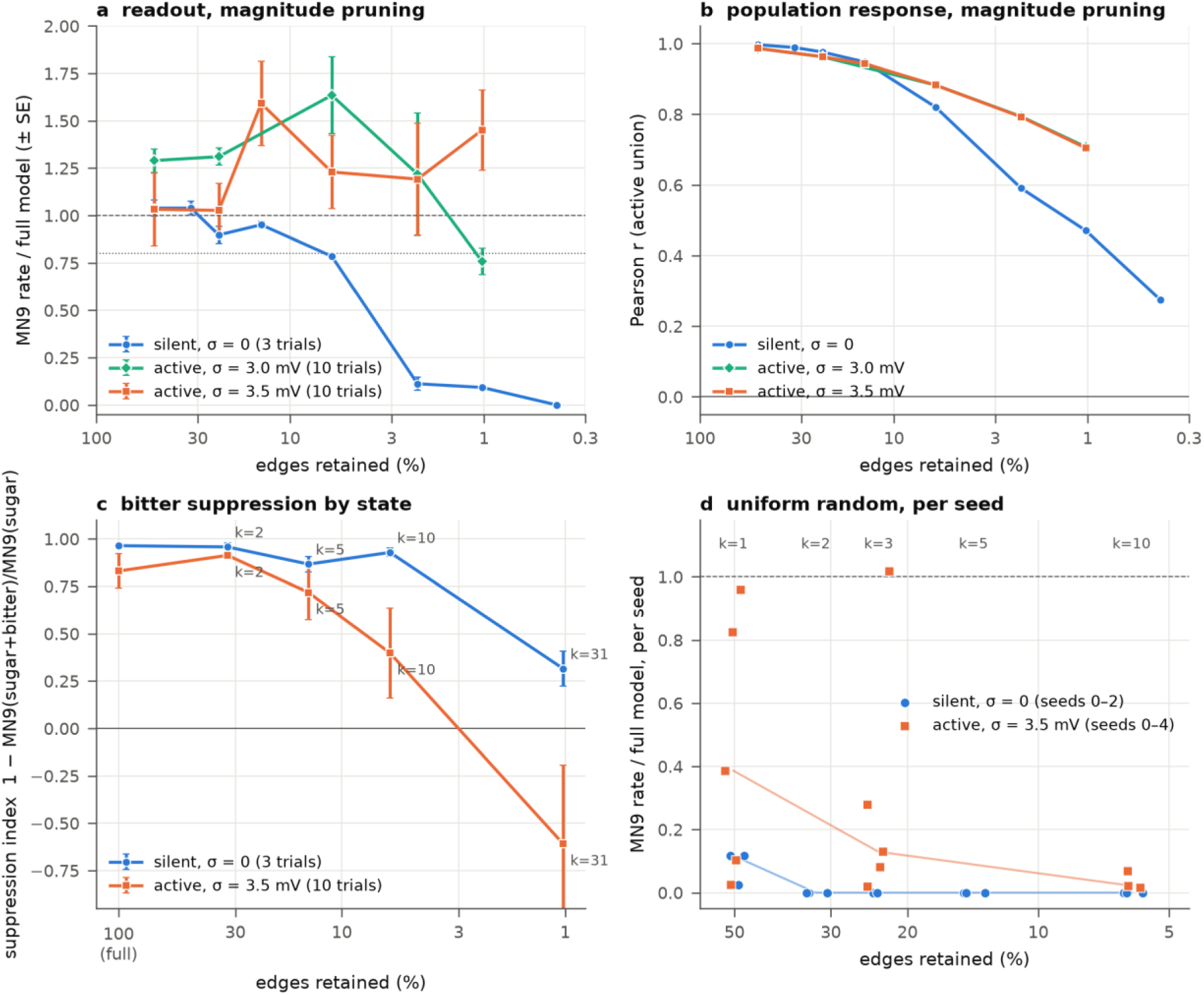
Network state changes how, not how much, sparsification fails. (a) Sugar readout ratio for magnitude pruning (± trial SE; σ = 0 3 trials, σ = 3.0/3.5 10 trials). In the active state the cliff does not move earlier; the readout exceeds the full model. (b) Population correlation declines gradually in all three states, with the σ = 3.0 and 3.5 curves superimposed. (c) Bitter suppression index SI = 1 − MN9(sugar+bitter)/MN9(sugar) in the silent (3 trials) and active (10 trials) states; error bars are delta-method SE. Suppression that holds at 0.93 up to k = 10 in the silent state declines from k = 5 in the active state to 0.40 at k = 10 and −0.61 at k = 31 (bitter now increases MN9). (d) Per-seed readout ratios of uniform random subsets. In the silent state (3 seeds) all are 0 for k ≥ 2; in the active state (5 seeds) they split bimodally between 0.02 and 1.02 at k = 1 and 3.

**Figure 5.**
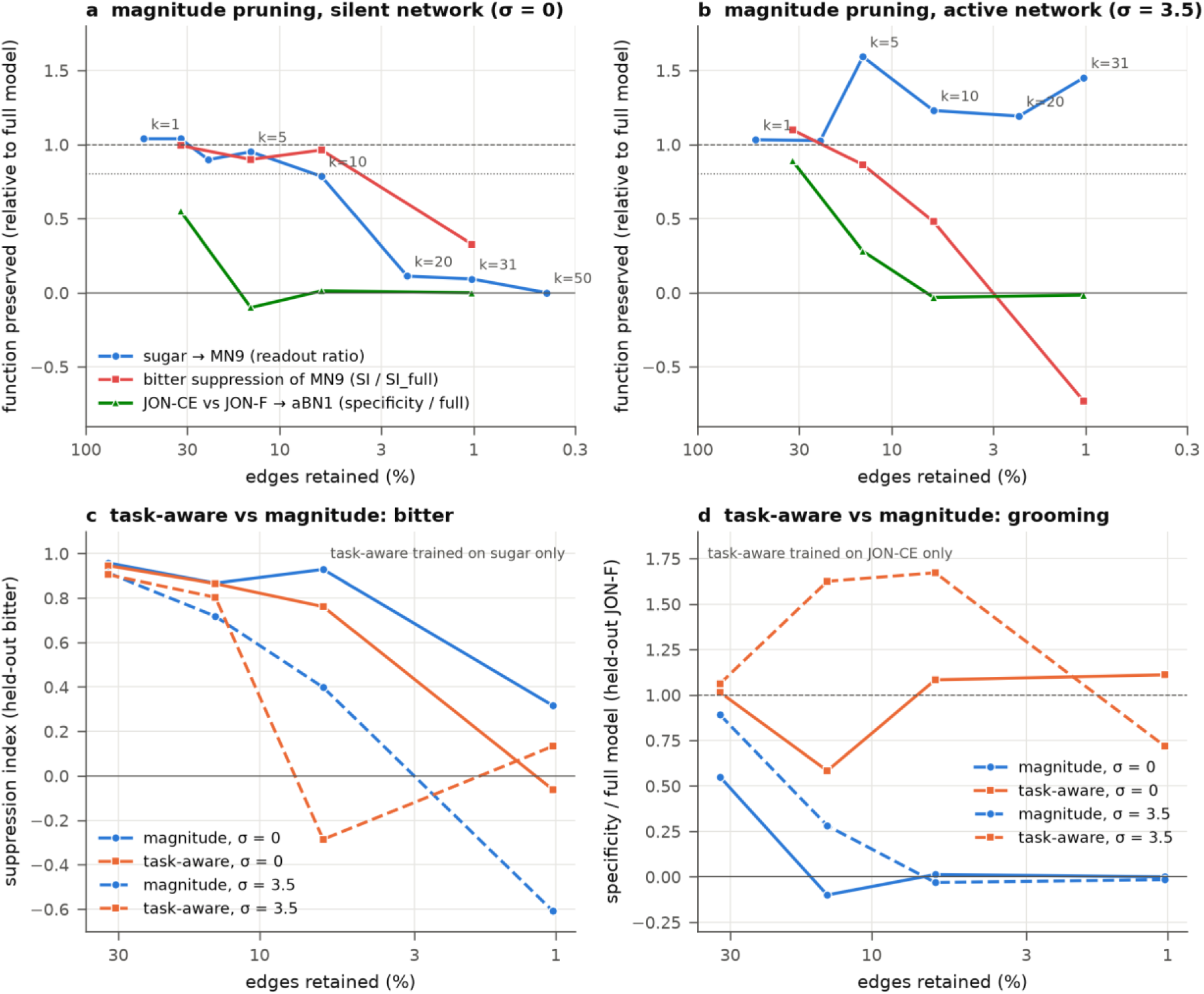
The cliff position depends on the task, and the task-aware criterion loses held-out functions selectively. (a, b) Preservation of three functions under magnitude pruning, relative to the full model: sugar → MN9 readout ratio (blue), bitter suppression index normalised by the full-model SI (red), grooming pathway specificity aBN1(JON-CE) − aBN1(JON-F) normalised by the full model (green). (a) Silent state; (b) active state (σ = 3.5; sugar and bitter from 10 trials). Grooming specificity is already halved at k = 2 and gone at k = 5, and in the active state bitter suppression collapses before sugar output. (c) Held-out bitter suppression index: the task-aware criterion trained on sugar only (orange) versus magnitude (blue), silent (solid) and active (dashed). (d) Held-out grooming specificity relative to the full model: the task-aware criterion trained on JON-CE only preserves or exceeds specificity, whereas magnitude pruning loses it at k = 5.

State dependence was clearest in **inhibitory function**. In the silent state the bitter suppression index held at 0.93 up to k = 10, but at σ = 3.5 (10 trials, Phase D) it declined progressively from 0.83 (full) to 0.91 at k = 2, **0.72 at k = 5, 0.40 at k = 10 and −0.61 at k = 31**, where inhibition had reversed into excitation. MN9 under sugar+bitter rose from 5.9 ± 2.9 Hz (full, mean ± SE) to 3.5 ± 0.7, 15.1 ± 7.3, 31.0 ± 11.1 and 48.1 ± 10.7 Hz, while sugar alone gave 34.8, 40.2, 53.5, 51.5 and 29.9 Hz. The 3-trial run (Phase B) showed the same direction more abruptly (0.92 at k = 5, 0.16 at k = 10, −0.17 at k = 31): the loss of inhibition is not a sudden collapse but a monotonic decline that begins at k = 5, ahead of the sugar cliff (k* ≈ 9). The grooming circuit collapsed at σ = 3.5 much as in the silent state (CE ratio 0.91, 0.46, 0.23 at k = 2, 5, 10; specificity 21.3 → 19.0, 6.0, −0.7). Uniform random controls, which die immediately in the silent state (0.02–0.12 at k = 1), were instead **all-or-nothing depending on the seed** in the active state: at k = 1 the five seeds gave ratios of 0.96, 0.39, 0.82, 0.02 and 0.10 (median 0.39) and at k = 3 0.13, 0.08, 1.02, 0.02 and 0.28 (median 0.13), a bimodal distribution, whereas for k ≥ 10 all were ≤ 0.07. With many neurons near threshold, whether a random edge set happens to connect the readout pathway decides whether output survives or dies—a threshold phenomenon not seen in the silent state.

In summary, network state changes the answer to "what goes wrong when we prune" more than the answer to "how much can be pruned": failure in the silent state is loss of output, failure in the active state is over-excitation and loss of inhibition. Numbers quoted for σ = 3.5 are from the 10-trial runs (sugar: Phases A2 and D; bitter: Phase D); the difference in reference values from Phases A and B (3 trials; 49.3 versus 37.0 Hz) lies within trial variability.

### 2.7 Behavioural output and neural response dissociate (H5)

Plotting the readout ratio against population correlation and Jaccard across budgets (Fig. 3), magnitude pruning keeps a readout of 0.95 at k = 5 while the active-set Jaccard is 0.83, and at k = 10 a readout of 0.79 with Jaccard 0.68 and 25% fewer active neurons—it enters a regime in which behaviour is preserved while the internal representation has already changed. In the active state the dissociation also appears in the opposite direction: the readout exceeds the full model while population correlation falls from 0.88 (k = 10) to 0.70 (k = 31). act, by contrast, preserves readout, correlation and Jaccard together on its training stimulus (1.06 / 0.997 / 0.92 at k = 50), so the dissociation is a property of the compression method, not a necessity of compression. That a compressed model can pass a behavioural test while already misrepresenting internal activity matters whenever a compressed connectome is used as a model of neural computation rather than as a controller.

### 2.8 What remains and what disappears

Weak-synapse pruning is sign-neutral in edge counts: the inhibitory fraction of removed edges is 0.39–0.40 at every k, equal to the whole graph (0.40; Fig. 6a). The inhibitory fraction of the retained synapse *mass*, however, rises from 0.40 at k ≤ 5 to 0.41 at k = 10, 0.43 at k = 20, 0.46 at k = 31 and 0.49 at k = 50—among strong edges, inhibitory ones are relatively more numerous. At the neuron level, the fraction of input-receiving neurons that lose *all* inhibitory input is 5.7% at k = 5, 17.6% at k = 10 and 28.8% at k = 20, and the fraction losing all excitatory input is 8.2%, 19.2% and 32.0%. The median inhibitory fraction of retained input stays at 0.44–0.46 (as in the full model) for k ≤ 10 and tilts to 0.51, 0.56 and 0.60 at k = 20, 31 and 50. The neighbourhood of k* (≈ 10) is thus the point at which about one fifth of neurons begin to lose one sign of input entirely; beyond it the remaining circuit is biased towards inhibition.

**Figure 6.**
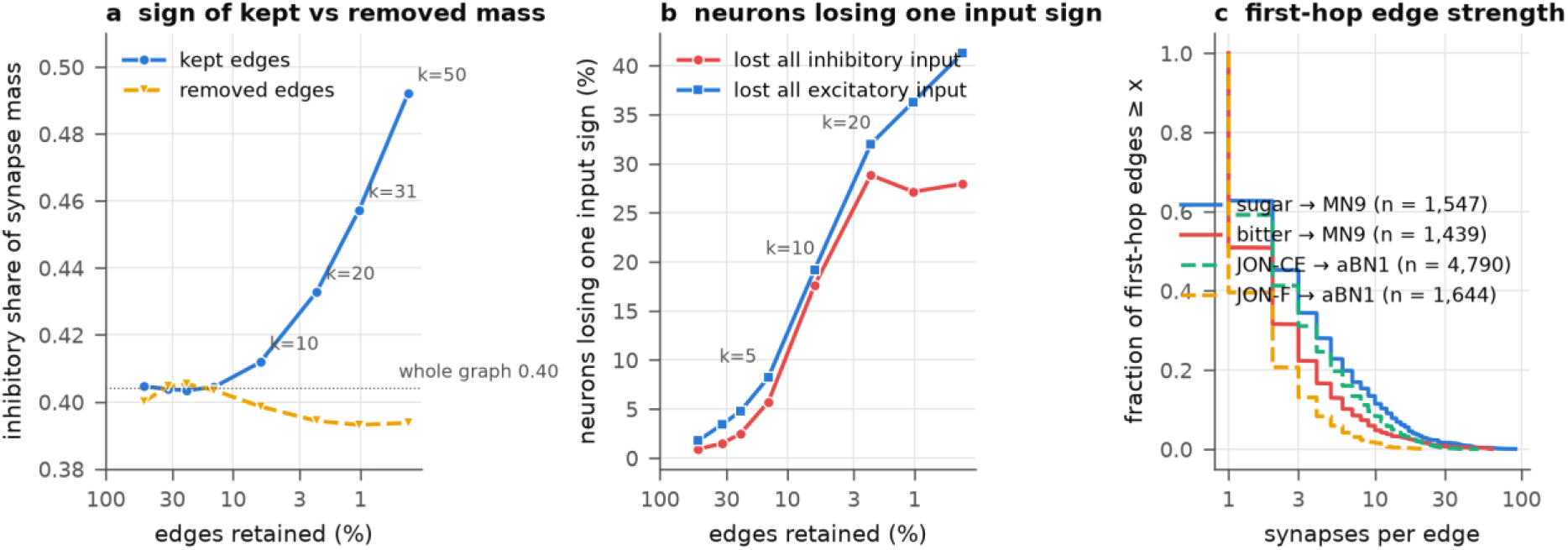
What remains and what disappears. (a) Inhibitory share of retained and removed synapse mass by budget. (b) Fraction of input-receiving neurons that lose all inhibitory or all excitatory input. (c) Synapse-count distribution of first-hop edges of the four validated pathways.

The first-hop edges of the four validated pathways are mostly weak: the fraction with ≤ 2 synapses is 55% for sugar → MN9, 68% for bitter → MN9, 59% for JON-CE → aBN1 and 79% for JON-F → aBN1, and the fraction with ≤ 5 synapses is 77–94%. That the sugar circuit nevertheless survives to k = 10 is due to the convergence of 21 sensory neurons and the redundancy of 286 intermediates. The JON-F pathway is the weakest of the four (median 1 synapse) and its input to aBN1 is inhibition-dominated (222 excitatory versus 671 inhibitory synapses), consistent with the disinhibition in §2.5 when weak inhibition goes first. By cell-class pair, at k = 2 70% of central → central edges are removed while only 46% of central → motor and 54% of ascending → motor edges are, so edges towards motor output are on average stronger and preserved. At the cliff k = 10 survival by super-class pair tilts towards output (Fig. S3): only 6.8% of central → central edges remain, versus 26.7% of central → motor, 21.7% of descending → motor, 15.8% of central → descending and 8.6% of sensory → central. The composition of the surviving graph is nevertheless dominated by the most numerous pairs, with central → central at 25% and optic → optic at 14%.

### 2.9 The cliff position depends on synaptic gain and, less, on connectome version

To ask whether the cliff at k* ≈ 9.4 is a property of the wiring or of the particular parameterisation, we repeated magnitude pruning at k = 5, 10 and 20 on the FlyWire v783 connectome (138,639 neurons, 15.1 M edges; 3 of the 174 stimulus and readout neurons are absent) and, on v630, with the synaptic gain *w_syn* scaled by 0.8 and 1.2 (Fig. S2; silent state, 3 trials). The full model’s sugar response was 65.0 Hz on v783 (v630: 68.3), 35.7 Hz at 0.8× gain and 81.0 Hz at 1.2× gain, so the gain alone moves the reference output by more than a factor of two. On v783 the readout ratio was 0.71 at k = 5, 0.70 at k = 10 and 0.06 at k = 20—a lower plateau than on v630 (0.95, 0.79, 0.11) but the same collapse between k = 10 and 20, with population correlations within 0.01 of the v630 values (0.94, 0.81, 0.60 versus 0.95, 0.82, 0.59). Gain moved the cliff itself: at 0.8× the ratio was already 0.22 at k = 5 and 0.06 at k = 10 (k* < 5), whereas at 1.2× it was 0.87, 0.97 and 0.56 (k* ≈ 13). The absolute position of the feeding-circuit cliff is therefore a property of the model’s operating point, not of the wiring alone, and the coincidence of k* ≈ 9.4 with the anatomical ≥ 10-synapse threshold holds only for the reference gain. The population-correlation curves, by contrast, were nearly invariant across all four conditions at k = 5 and 10 (0.92– 0.98 and 0.81–0.93), and behaviour was lost before the neural response in every condition (at 0.8× gain the correlation was still 0.86 when the readout had fallen to 0.06)—the dissociation of §2.7 is robust where the cliff position is not.

## 3. Discussion

### Summary

In an experimentally validated fly whole-brain model, weak-synapse pruning preserves sugar-evoked behaviour after removing 94% of edges, but this limit (i) holds only when placement and strength are kept together, (ii) moves much earlier when the circuit requires inhibition or pathway specificity, and (iii) changes its mode of failure from silencing to over-excitation when the network is spontaneously active. The limit of function-preserving compression is not a fixed property of the graph but a function of graph, task and state.

### Relation to anatomical thresholds

The synapse-count thresholds that connectome modelling inherited from anatomy (≥ 5, ≥ 10; our grid points k = 5 and 10 are ≥ 6 and ≥ 11, immediately adjacent) lie *inside* the function-preserving region for the sugar circuit (k = 5: readout 0.95) or at its boundary (k = 10: 0.79). For grooming pathway specificity, however, half is already lost at k = 2 and all at k = 5, so the same thresholds lie *outside*. The first practical implication is that the choice implicitly made by FLYNN, effectome analysis and neuromorphic mapping is safe for feedforward output but not for inhibition-dependent function. That Schlegel’s 90% reproducibility point (k = 10) coincides with the sugar cliff at k* ≈ 9.4 would be consistent with the hypothesis that "evolutionarily conserved connections = functionally required connections", but the coincidence is fragile: a 20% change in synaptic gain moves the cliff to below k = 5 or to k ≈ 13 (§2.9), and the grooming circuit depends on much weaker connections. The anatomical threshold and the functional cliff should therefore not be equated; what is robust across gain and connectome version is the ordering of the operators and the dissociation between behaviour and population response.

### Why degree statistics are not enough

Our degree-preserving null differs from Dhiman’s in an important respect: it is applied to a fixed, untrained spiking model, so rewiring cannot be compensated by learning. The collapse we observe therefore isolates the contribution of specific wiring, while the disappearance of the advantage reported in trained networks reflects what learning can restore. The two results are consistent: wiring specificity matters in a fixed model and can be partly relearned in a trainable one. That weight shuffling also dies—as loss of output in the feeding circuit and as a seed-dependent lottery over which pathway drives the output in the grooming circuit—means that topology (where connections go) is not sufficient either, and quantifies what approaches that use the connectome as a binary adjacency matrix (some FLYNN variants) give up. The weight-proportional random control adds a third argument. The simplest alternative explanation, "random controls die because they lose synaptic mass", does not hold: the mass retained by randm declines gently from 92% (k = 1) to 72% (k = 5) of the magnitude criterion’s, yet function drops abruptly between k = 3 and 5, and two graphs of equal mass (rand k = 1 and randm k = 3, both ≈ 50%) give 0.09 and 0.90. Function is carried not by a total but by the integrity of a specific edge set, which predicts that compression rules that miss strong edges even stochastically (stochastic sparsification, thresholding after quantisation) will do worse than a deterministic magnitude criterion at the same budget.

### Inhibition dies first

Three results point the same way: grooming specificity collapses through disinhibition, the task-aware criterion discards inhibitory pathways first, and in the active state bitter suppression declines from k = 5, falls below half at k = 10 and reverses at k = 31. Anatomically, weak-synapse pruning is sign-neutral in counts, yet at k = 10 18% of neurons lose all inhibitory input. Inhibition tends to be implemented by more *distributed* weak connections than excitation (the 671 inhibitory synapses from JON-F intermediates onto aBN1 are spread over 78 edges) and is therefore vulnerable to any per-edge removal rule. This suggests a design principle for the next step: compression criteria should constrain per-neuron E/I balance rather than act edge by edge.

### State dependence

Pre-registered prediction A (an earlier cliff in the active state) was not supported by the readout ratio; instead the mode of failure changed. Two interpretations are possible. First, the function-preserving limit of the silent LIF model is not an overestimate for the real brain but misses *a different kind of risk*: over-excitation through loss of inhibition. Second, the spontaneous activity produced by our noise model (uniform white noise) is heavy-tailed (p99 77 Hz) and includes recurrent amplification in high fan-in neurons, so part of the over-excitation may be an artefact of the noise model. The near-identical population-correlation curves at σ = 3.0 and 3.5 show robustness to noise amplitude. Trial variability of the readout is large (at σ = 3.5 the SD is 69% of the full-model mean), but the monotonic decline of the suppression index measured with 10 trials and the bimodal distribution of the random control lie outside that variability. The all-or-nothing behaviour of uniform random subsets in the active state reads as the same mechanism: with many neurons near threshold, an arbitrary subgraph has a non-zero probability of connecting the readout pathway; when it does, background activity amplifies the pathway, and when it does not, the network is as silent as in the resting state. This threshold phenomenon also means that a compressed graph should never be evaluated from a single random realisation.

### Behaviour is a lenient test

Magnitude pruning at k = 5–10 keeps behaviour while changing 17–32% of the active set. Evaluations of compressed or hardware-mapped models that report only output rates (F. Wang et al., 2025) can pass models whose internal representation has already shifted. That act preserves both on its training stimulus shows the dissociation is not inevitable, but the price is the transfer failure of §2.4.

### Implications for pruning artificial networks

Magnitude pruning is the default rule in artificial-network compression (Han et al., 2015), and these results suggest why a rule of that family eventually fails. What survives is not the set of large weights as such, but the intersection of large weights with particular wiring positions: weight-proportional random sampling retains most of the synaptic mass and still collapses at the budget where it begins to miss individual strong edges (§2.3). If this carries over to trained networks—which our fixed model cannot establish, only motivate—then rules that retain strong weights in expectation rather than deterministically (stochastic sparsification, thresholding after quantisation) should be at a disadvantage at equal budget, and the weight-proportional control used here measures the size of that disadvantage in a setting where retraining cannot mask it. It also suggests a reading of the recovery step that dominates the artificial-network literature (Gale et al., 2019; Renda et al., 2020): retraining may be doing less repair of a damaged function than re-derivation of a new one, a difference that is invisible whenever the only score is task accuracy. The dissociation of §2.7 adds a second caution: a compressed network that matches its teacher on the task metric may already differ in internal representation, which matters whenever the compressed model is reused for another task (§2.4) or interpreted as a model of the computation rather than as a controller.

### Relation to concurrent work

Three recent studies ask neighbouring questions with different designs, and the comparison sharpens what is specific here. Dhiman (2026) and McAllister et al. (2026) both *train* networks whose connectivity is taken or initialised from a fly connectome and compare them against sparsity-or degree-matched random wiring; the former reports that the topological advantage claimed for connectome initialisation largely disappears under a fair null, the latter that connectome wiring supports robust and efficient function at high sparsity across a battery of cognitive tasks. Therianos (2026) applies a frozen rate operator to the complete larval connectome with a degree-and-weight-matched rewiring ensemble and reports a division of labour close to ours: degree and weight govern the gross response, exact wiring governs routing. Our design differs from all three in that the teacher is a spiking model whose input–output behaviour has been validated experimentally and whose parameters are never fitted, so the quantity preserved is a behaviour rather than a task accuracy and no recovery step can mask what removal costs; and it is the edge budget, rather than the wiring rule, that is held fixed across conditions. That a similar separation between bulk response and specific routing appears in trained networks, in a frozen rate model of the larva and in a validated spiking model of the adult suggests it is a property of connectome graphs rather than of any one simulator.

### Limitations

(1) All conclusions concern one validated *model* of the fly brain—one connectome version (v630) and one parameter set. The LIF model omits neuromodulation, electrical synapses, plasticity and morphology, and assumes zero baseline firing at rest. A ± 20% change in synaptic gain moves the feeding-circuit cliff between k < 5 and k ≈ 13 (§2.9), so absolute cliff positions reported here should be read as properties of the reference operating point; the operator ordering and the behaviour–neural dissociation were robust to gain and to the v783 connectome, but the null models and the other tasks were tested only at the reference gain on v630. (2) Function is defined by three circuits of the gustatory and mechanosensory systems; visual and navigational functions were not tested. (3) The task-aware criterion is deliberately simple (activity-weighted magnitude); gradient-based saliency in a differentiable model is the natural next step. (4) The active state uses uniform white noise, and the heavy-tailed firing of high fan-in neurons is a consequence (S1). (5) The pre-registered 5 seeds and 5 (or 10) trials were met only for the key conditions (σ = 3.5 sugar and bitter magnitude pruning; uniform and weight-proportional random at k ≤ 5); the rest use 3 seeds and 3 trials. Individual trial values were not stored, so we report SE and seed ranges rather than trial-level bootstraps. (6) The water circuit was omitted for lack of time.

### Outlook

Function-preserving sparsification is a common benchmark for the growing family of connectome-constrained models, and a route to hardware-and memory-efficient whole-brain simulation that measures fidelity instead of assuming it, in the tradition of spiking models built to compute rather than only to describe (Maass, 1997; Tavanaei et al., 2019). The finding that inhibition goes first motivates compression criteria with per-neuron E/I constraints; the transfer failure across tasks motivates decomposing a shared core trained on several stimuli; and the male central nervous system connectome (Berg et al., 2026) motivates comparing compression limits between sexes and individuals.

## 4. Methods

### 4.1 Connectome and reference model

We use the connectivity table released with Shiu et al. (FlyWire materialisation 630: 127,400 proofread neurons, 14,687,178 edges; each edge *w_ij* is the synapse count from *i* to *j*, multiplied by −1 if *i* is predicted GABAergic or glutamatergic). Neurons are single-compartment LIF units with alpha synapses: *τₘ v̇ = (v_0 - v) + g*, *τₛ ġ = -g*, *v_0=v_reset=-52* mV, *v_th=-45* mV, *τₘ=20* ms, *τₛ=5* ms, refractory period 2.2 ms, synaptic delay 1.8 ms; a presynaptic spike increments *g_j* by *w_ij· w_syn* (*w_syn=0.275* mV). All parameters are those of Shiu et al. and were not tuned. Simulations use Brian2 (v2.10; Stimberg et al., 2019) with a 0.1 ms time step; the silent condition uses the exact linear integrator and the noisy conditions Euler integration.

### 4.2 Stimulus protocols and readouts

Following Shiu et al., stimulated neurons receive independent Poisson input of weight *250 w_syn* at rate *r*. Protocols and validated readouts: (i) *sugar*: 21 right-labellar sugar GRNs at 100 Hz, readout MN9 (proboscis extension); (ii) *bitter*: sugar GRNs at 100 Hz plus 21 bitter GRNs at 100 Hz, readout MN9, suppression index *SI = 1 - MN9_sugar+bitter/MN9_sugar*; (iii) *groom*: 70 JO-CE neurons at 100 Hz and, separately, 60 JO-F neurons at 100 Hz, readout aBN1, specificity *= aBN1_CE - aBN1_F*. Neuron IDs are those released with Shiu et al. Each condition is simulated for 1 s from rest and repeated over *n_trial* independent Poisson realisations.

### 4.3 Sparsification operators

Let *b_k = |{(i,j): |w_ij|>k}|*. *Magnitude (mag)*: keep exactly those edges. *Uniform random (rand)*: a uniform subset of size *b_k*. *Weight-proportional random (randm)*: a subset of size *b_k* drawn without replacement with probability proportional to *|w_ij|*—a stochastic control that approaches the synaptic mass of mag*_k* without a deterministic threshold; it retains 92, 85, 80, 72 and 58% of mag*_k*’s mass at k = 1, 2, 3, 5, 10 and overlaps mag*_k*’s edge set by 71, 61, 54, 46 and 34% (Table S2). *Degree-preserving (dp)*: apply a uniform permutation to the postsynaptic index column of the magnitude set, preserving every neuron’s out-and in-degree and the sign of every retained edge (Dale’s law) while destroying placement; this is the degree-preserving randomisation standard in network neuroscience (Maslov & Sneppen, 2002; Rubinov & Sporns, 2010), applied to the thresholded graph so that its budget matches mag*_k* exactly. *Weight shuffle (shuf)*: permute *|w_ij|* among the edges of the magnitude set, keeping signs; preserves placement and destroys magnitude. *Task-aware (act)*: *s_ij=|w_ij|(1+r_i)*, where *r_i* is the mean rate of neuron *i* in the full model under the task’s first stimulus protocol (the training stimulus; sugar100 or jonCE100); keep the top *b_k*. The second stimulus (bitter, jonF) is held out. Random operators are run with seeds 0–2 (0–4 in Phase D); deterministic operators once.

### 4.4 Network state

Spontaneous activity is induced by additive membrane noise: *τₘ v̇ = (v_0 - v) + g + σ√τₘ ξ(t)*, with *ξ* unit white noise. *σ* was calibrated in the full model over 300 ms: *σ=2.5* mV produces almost no spontaneous spikes (0.7% of neurons active), *σ=3.5* mV a mean of 4.3 Hz with 25.5% of neurons active (median 0, 90th percentile 6.7 Hz, 99th percentile 77 Hz). In the 1 s stimulated simulations 78–93% of neurons fire at least once at both σ = 3.0 and 3.5. We use *σ ∈ {0, 3.0, 3.5}* mV.

### 4.5 Metrics

For each stimulus, per-neuron mean rates *r̄ᶠᵘˡˡ* and *r̄’* are computed over trials; stimulated neurons are excluded from all population metrics. *Readout ratio = r̄’_out/r̄ᶠᵘˡˡ_out*. *Active set A = {i: r̄_i > 0}*; *Jaccard = |Aᶠᵘˡˡ∩ A’|/|Aᶠᵘˡˡ∪ A’|*. *Population correlation* = Pearson *r* between *r̄ᶠᵘˡˡ* and *r̄’* over *Aᶠᵘˡˡ∪ A’* (undefined, and reported as collapse, if the union has fewer than 3 neurons or zero variance). *Cliff k\**: the budget at which the readout ratio first falls below 0.8, log-interpolated between tested *k*. *Dissociation index* = readout ratio − population correlation. Suppression index and specificity as in §4.2.

### 4.6 Anatomical analysis

Using FlyWire annotations (super_class, cell_type) we tally removed and retained edges at each *k* by sign and by cell-class pair, and compute for each input-receiving neuron the inhibitory fraction of its retained input and the fraction of neurons losing one sign entirely. For the four validated pathways we tally the synapse-count distribution of first-hop edges leaving the sensory neurons and the excitatory and inhibitory mass from their postsynaptic partners (intermediates) onto the readout neuron.

### 4.7 Statistics

Silent state: random operators 3 seeds × 3 trials (randm at k = 3, 5: 5 seeds), deterministic operators 3 trials. Active state (σ = 3.5), numbers quoted in the text: sugar deterministic operators and rand 10 trials (rand at k = 1, 3: 5 seeds), bitter magnitude pruning 10 trials, everything else (groom; dp, shuf, act) 3 trials. The across-trial SD of the readout (population SD, ddof = 0) is stored for every condition; uncertainty for deterministic operators is reported as SE = SD/√n and for random operators as per-seed values with median and range (individual trial values were not stored, so trial-level bootstraps were not performed). The full-model reference was re-simulated in each run unit (Phase), so reference values for the same condition differ between Phases by trial variability (e.g. grooming specificity 26.7 Hz in Phase B versus 24.0 Hz in Phase C; σ = 3.5 sugar 49.3 Hz with 3 trials versus 37.0 Hz with 10); ratio metrics are always computed against the reference of the same Phase. All conditions of the pre-registered design (S1) are reported regardless of outcome. The pre-registered target of 5 seeds and 5 trials was met for the key conditions (above); the rest use 3 seeds and 3 trials. Robustness runs (§2.9; v783, gain 0.8× and 1.2×) use magnitude pruning only, 3 trials, with their own full-model reference; the runner options -- connectome 783 and --w-syn-scale reproduce them.

### 4.8 Computational resources

Silent-state simulations ran on a laptop CPU (Intel i3-1315U, 8 GB RAM); the noisy conditions and Phases B and C ran on a Google Colab CPU runtime, all with the Brian2 NumPy backend. One whole-brain 1 s trial takes ≈ 65 s in the silent condition and ≈ 100–180 s with noise. Code, neuron lists, the pre-registered design and every summary table are released in the repository of §4.9.

### 4.9 Data and code availability

The runner (run_flylite.py), stimulus and readout neuron IDs (task_ids.json), analysis and figure scripts, the pre-registered design document, the per-condition summary tables of Phases A–D and F (results/phase*_summary.csv, 371 rows in total), the randm–mag overlap table, the anatomical tallies and the figure sources are released under the MIT licence at https://github.com/aikian/flylite. The reference model code and connectome data are obtained from the repository of Shiu et al. (github.com/philshiu/Drosophila_brain_model). Every number quoted in the manuscript is cross-checked against the tables by scripts/check_numbers.py.

## Figure legends

**Figure S1.**
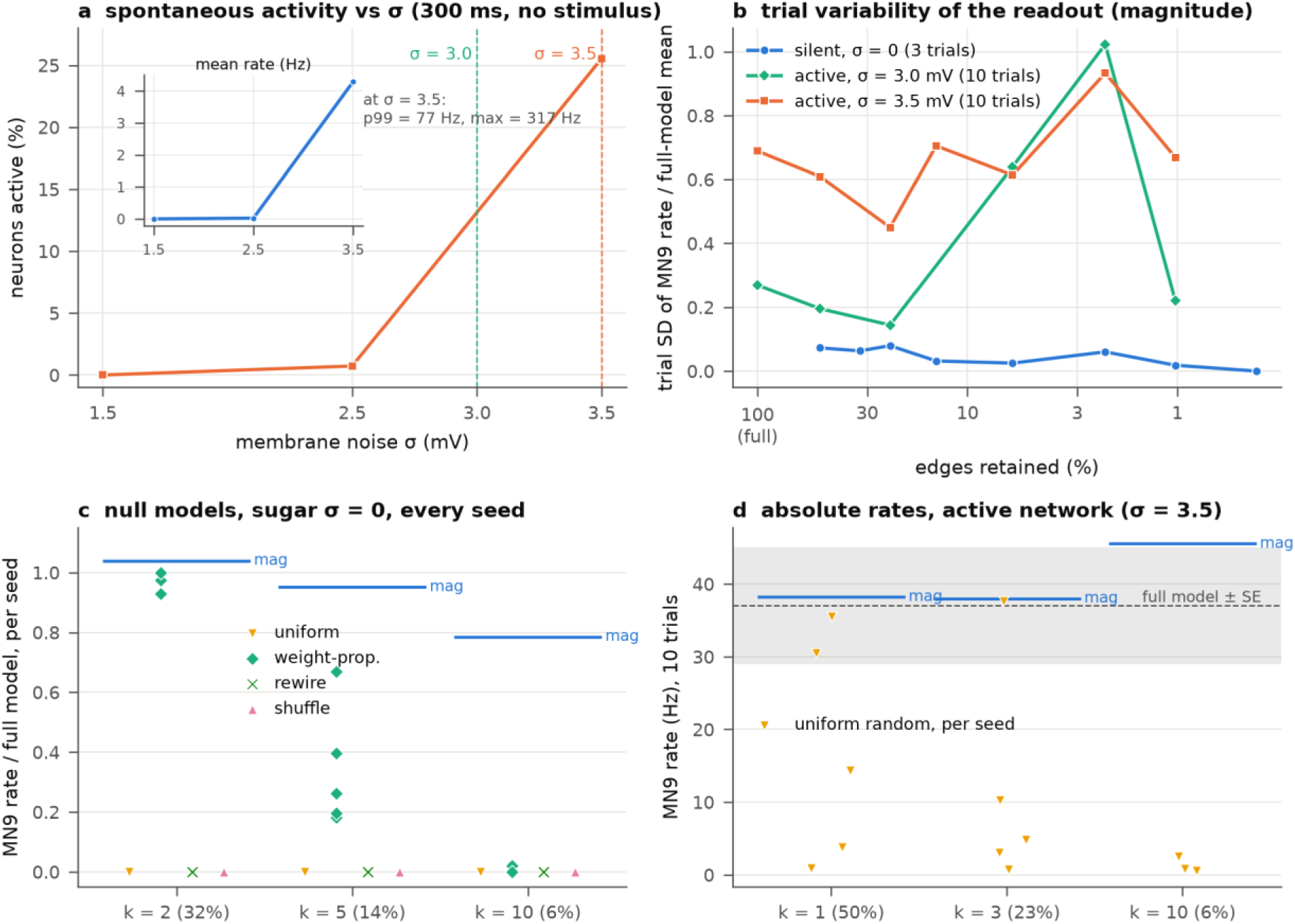
Calibration and variability. (a) Spontaneous activity of the full model as a function of membrane noise σ (no stimulus, 300 ms): fraction of active neurons (main axis) and mean rate (inset). The network switches on where the 7 mV threshold is 2–3 times the steady-state SD (σ/√2); at σ = 3.5 the distribution is heavy-tailed (p99 = 77 Hz). (b) Across-trial SD of the readout under magnitude pruning, normalised by the full-model mean, by state. In the active state the SD reaches 45–100% of the mean, which is why 10 trials were added. (c) Per-seed readout ratios of the four null models in the silent sugar task (k = 2, 5, 10; blue lines = magnitude pruning). (d) Absolute MN9 rates in the active state (σ = 3.5, 10 trials): uniform random per seed (yellow), magnitude pruning (blue lines), full model ± SE (grey band). The bimodal distribution of uniform random lies outside the uncertainty of the reference.

**Figure S2.**
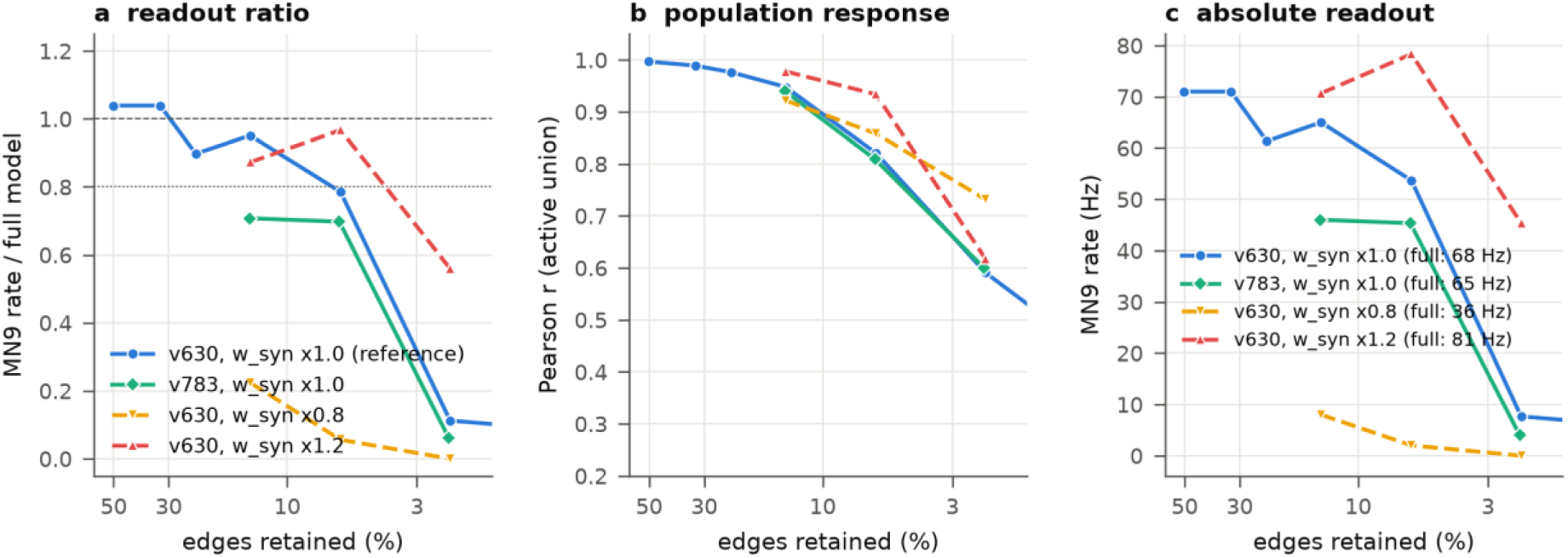
Robustness of the feeding-circuit cliff to connectome version and synaptic gain. Magnitude pruning at k = 5, 10, 20 (silent state, 3 trials) on the FlyWire v783 connectome and on v630 with the synaptic gain *w_syn* scaled by 0.8 and 1.2, against the v630 reference of Fig. 1b. (a) Readout ratio relative to each condition’s own full model. (b) Population Pearson correlation. (c) Absolute MN9 rate; full-model rates in the legend. Gain shifts the cliff (0.8×: k* < 5; 1.2×: k* ≈ 13) while the correlation curves and the behaviour-before-neural dissociation are preserved; v783 lowers the plateau (0.70 at k = 5–10) but collapses at the same budget.

**Figure S3.**
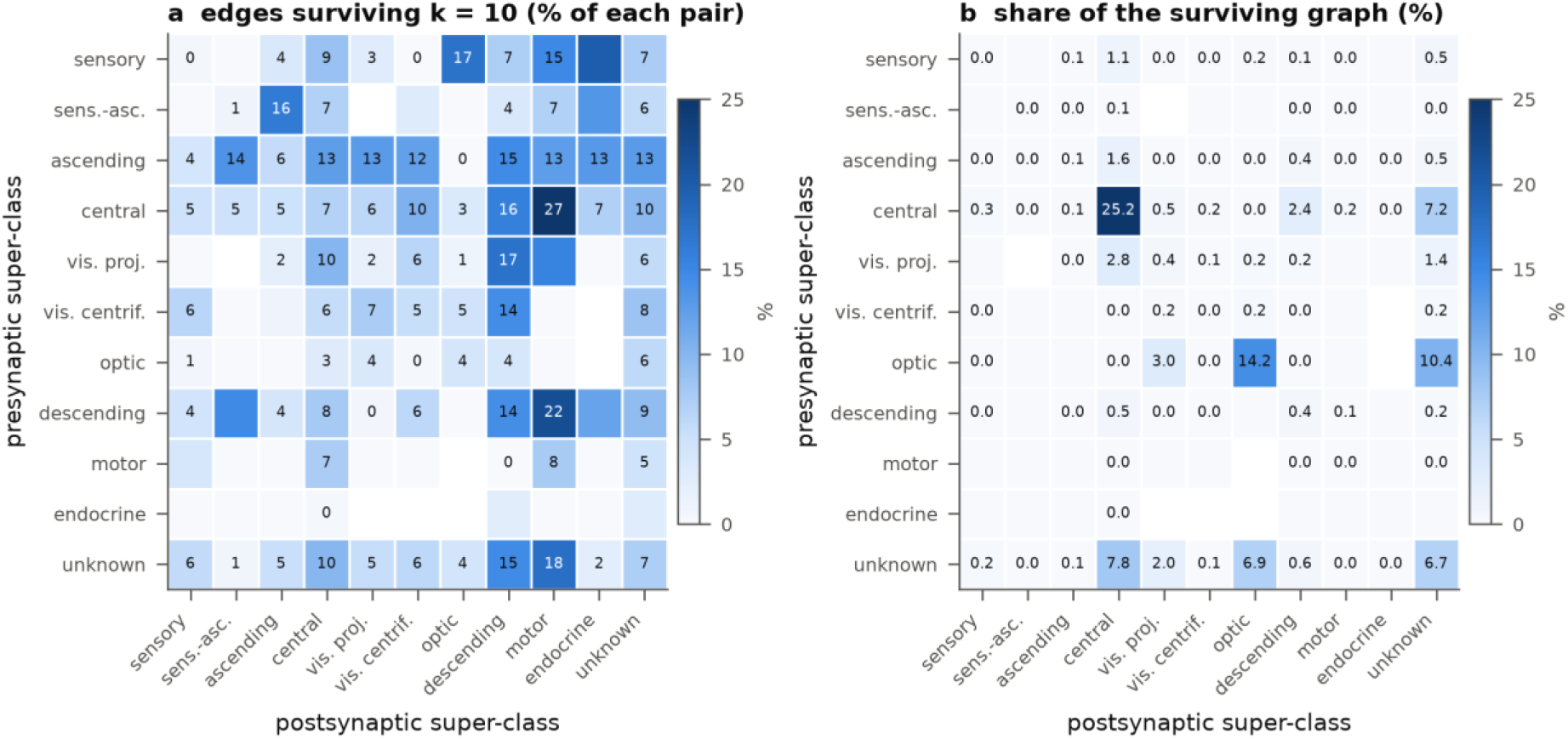
What remains at the cliff. Per super-class pair (rows = presynaptic, columns = postsynaptic) at magnitude pruning k = 10 (6.1% of edges): (a) fraction of edges retained (%) and (b) share of the surviving graph (%). Numbers are omitted for pairs with fewer than 200 edges. Edges onto motor and descending neurons survive 2–4 times more often than central-internal edges.

**Table S2.** Overlap between weight-proportional random (randm) and magnitude (mag). Computed by reproducing randm’s edge sets with the same seeds (analyze_phaseC.py).

| k | edges | mass: mag | mass: randm | mag_k edges included in randm | mag_k mass included in randm | weak ($\leq k$) edges in randm | edges > 10 synapses retained | edges > 31 synapses retained |
| --- | --- | --- | --- | --- | --- | --- | --- | --- |
| 1 | 50.3% | 86.2% | 78.9% | 71% | 87% | 29% | 99% | 100% |
| 2 | 32.5% | 76.3% | 65.0% | 61% | 79% | 39% | 93% | 100% |
| 3 | 23.3% | 68.6% | 54.9% | 54% | 72% | 46% | 83% | 99% |
| 5 | 14.1% | 57.3% | 41.2% | 46% | 63% | 54% | 63% | 91% |
| 10 | 6.1% | 40.7% | 23.7% | 34% | 47% | 66% | 34% | 63% |

**Supplementary Note S1.** Pre-registered hypotheses and verdicts.

| Hypothesis | Pre-registered prediction | Result | Verdict |
| --- | --- | --- | --- |
| H1 cliff | held to $k \leq 10$, sharp drop by $k \geq 31$ | $k^* \approx 9.4$ (sugar, $\sigma = 0$); 0.11 at $k = 20$ | as predicted; coincides with Schlegel's 90% point |
| H2 state | A: earlier cliff / B: unchanged | readout over-excites ($> 1$), correlation declines gradually; SI (10 trials) 0.83 → 0.91, 0.72, 0.40, -0.61 ($k$ 2, 5, 10, 31); uniform random all-or-nothing by seed ($k = 1$: 0.96, | neither A nor B → reinterpreted as "a change in the mode of failure"; inhibition declines monotonically from $k = 5$ |
|  |  | 0.39, 0.82, 0.02, 0.10) |  |
| H3 placement · strength | dp dies, shuf partially preserved | dp 0 (all conditions); shuf: sugar and bitter 0 ($k \leq 10$, $\sigma$ 0 and 3.5), grooming fluctuates 0.03–2.4 by seed with random specificity; randm: held for $k \leq 2$ (0.93–1.01), mean 0.86 at $k = 3$ (5 seeds, 0.64–0.97), collapse at $k = 5$ (0.34 vs mag 0.95), 0.02 at $k = 10$—at equal mass (41%) mag $k = 10$ gives 0.79 | interpretation-table row 2: both are needed; the "random controls lose mass" objection is rejected |
| H4 task | inhibition and specificity collapse first | grooming $k^* < 2 \ll$ sugar 9.4; bitter SI holds to $k = 10$ at $\sigma = 0$, lost from $k = 5$ at $\sigma = 3.5$ | grooming as predicted; bitter more robust than expected at rest (collapses first, as predicted, in the active state) |
| H5 dissociation | readout and correlation curves separate, more so with noise | $k = 5$: 0.95 vs Jaccard 0.83; opposite-direction dissociation with noise | present; "more so" undecidable because of readout variability |
| H6 task-aware transfer | training act ≥ mag, held-out act ≤ mag | training: 1.06 even at $k = 50$; held-out SI 0.76 vs 0.93 at $k = 10$, $-0.06$ at $k = 31$; held-out grooming (jonF) specificity preserved (26.7 vs 24.0 at $k = 31$) | bitter as predicted, grooming transfers → "absence is free, presence is lost" |

## Declaration of generative AI and AI-assisted technologies in the writing process

During the preparation of this work the author used Claude Opus 5 (Anthropic), through the Claude Code command-line interface, to draft and edit the text of this manuscript, to write the simulation, analysis and figure-generation scripts released with it, and to cross-check the reported numbers against the result tables. After using this tool, the author reviewed and edited the content as needed and takes full responsibility for the content of the published article.

